# Slowed aquatic invasion coincides with intensified research and prevention in Minnesota, USA

**DOI:** 10.64898/2026.11.06.764903

**Authors:** Jake R. Walsh, Alex W. Bajcz, Amy C. Kinsley, Nicholas B. D. Phelps

## Abstract

Prevention is often the most cost-effective expenditure of limited environmental management resources. While demonstrating that harms occur without prevention is relatively straightforward, showing harms have lessened due to prevention—proving its effectiveness—is not, in part because it requires counterfactual data (i.e., quantification of harms that did not occur and then attributing these to prevention). Here, we attempt to measure aquatic invasive species (AIS) prevention effectiveness in Minnesota, USA, where AIS, their management, and research have increased since the 1980s. We showed the rate of new aquatic invasions slowed alongside intensified prevention investment starting around 2015. Further, mechanistic simulations grounded in real boater movement and boat inspection effectiveness data suggested prevention may have reduced new invasions of two high-priority AIS by 20-53% over a ten-year span. We argue our evidence strongly supports long-term statewide prevention, monitoring, and research.

## Introduction

Managers are tasked with evaluating different prevention strategies to effectively and defensibly manage ecosystems given limited resources (Moore et al. 2021). A challenge to these efforts is the lack of data and guidelines with which to assess policy and program outcomes, which is magnified when the desired outcome is preventing adverse outcomes (Ferraro and Pattanayak 2006). Invasive species prevention exemplifies these challenges (Chivers et al. 2017). Invasive species have been associated with massive ecological and economic losses (Diagne et al. 2021), especially in aquatic systems (Moorhouse and Macdonald 2015). Thus, protecting ecosystems and economies through successful aquatic invasive species (AIS) prevention is thought to be cost-effective (Vander Zanden and Olden 2008; Epanchin-Niell and Liebhold 2015). However, this assertion hinges on being able to objectively measure prevention effectiveness, which, despite emerging evaluation tools, remains challenging due to the complexity of interacting invasion dynamics, management, and policy (Epanchin-Niell 2017).

Integrative approaches have proven successful in evaluating prevention effectiveness despite system complexity (Ferraro et al. 2015). Epidemic theory offers a particularly useful conceptual foundation for an integrative approach that also closely parallels invasion theory (Hulme et al. 2020). Changes in disease transmission intensity are commonly used to assess the effectiveness of prevention in public health and epidemiology (Anderson and May 1991). When empirical data are limited or when alternative intervention scenarios are needed, simulation models are often used to estimate transmission intensity and mimic the effects of prevention under plausible assumptions (Keeling and Rohani 2008). Similar modeling frameworks have been applied to AIS spread, offering analogous insight into how interventions alter invasion dynamics (Ogden et al. 2019).

### Leveraging a history of AIS prevention investment

Minnesota’s history of AIS research and management provides a unique opportunity to study AIS prevention effectiveness. The Minnesota Department of Natural Resources (MNDNR) has documented annual statewide AIS management activities, spending, and spread date back to 1988. Funding for AIS management in Minnesota increased from ∼$4 million/year from 1988-2008 to ∼$9 million/year in 2014 (Appendix S1: Figure S1). In 2015, the State augmented this funding via an additional appropriation of $10 million/year to counties, which is expected to be reduced to $5 million/year in 2027 (Minnesota Statute 4771A.19 Subd. 5). Complementary funding at local, tribal, or lake levels provide additional support for prevention that likely increased along with State funding (e.g., Ibrahim et al. 2017).

Minnesota has also benefited from strong research investments, largely funded by the State’s Environment and Natural Resources Trust Fund. The Minnesota Aquatic Invasive Species Research Center at the University of Minnesota (MAISRC) was established in 2012 and has worked with partners to develop research-based solutions to AIS problems. For example, MAISRC’s “AIS Explorer” web application simulates the movement of AIS between waterbodies under contrasting management scenarios (Kinsley et al. 2024; N. R. Angell et al. 2025). The underlying models are grounded in decades of AIS movement and management data documented by State and local agencies and organizations, allowing for development of “what-if” scenarios – a powerful tool that can be used for approximating counterfactual rates of invasion.

We leveraged this unique management and research history to evaluate the effectiveness of Minnesota’s AIS prevention. We combined State data documenting AIS spread with simulation and analytical modeling to 1) confirm that Minnesota lakes are unlikely to be saturated with AIS, 2) explore how rates of spread map to rates of prevention investment, and 3) estimate how many invasions might otherwise have occurred without that investment. We demonstrate that long-term commitment to invasive species management, documentation, and research can yield the ever-elusive defensible evidence that prevention can be effective. Not only does this provide support for ongoing investment in prevention, to our knowledge, this evidence has not been presented at such a broad scale. We argue that Minnesota offers rare evidence of and a valuable model for prevention success.

## Methods

All data manipulation, analyses, and visualizations were conducted using the statistical program R (R Core Team 2026).

### Invasion Records

Invasion records were obtained from the Minnesota Department of Natural Resources’ (MNDNR) Infested Waters List (Minnesota Department of Natural Resources 2025a). Each record includes the year the waterbody was designated infested by one of 12 high priority AIS (Appendix S1: Table S1). We only considered waterbodies for which the AIS was confirmed by physical specimen identification with a year of infestation.

### Estimating AIS saturation in Minnesota lakes: Is prevention still warranted?

With many AIS present in Minnesota for decades, one may question the premise that there are still potential invasions to prevent. Using state invasion records, we first determined how many lakes were designated as infested with each of the ten most prominent AIS. We then estimated how many invasions were theoretically left to prevent under a range of potential suitability assumptions (what fraction of lakes can be invaded?) and true invasion rate (what fraction of current invasions are documented?). We based our lake suitability estimates on past studies suggesting between 30% and 70% of Minnesota’s lakes are suitable for starry stonewort (*Nitellopsis obtusa*), zebra mussel (*Dreissena polymorpha*), and Eurasian watermilfoil (*Myriophyllum spicatum*) (Muthukrishnan et al. 2018; Kanankege 2019), which account for 759 of 894 invasions on the Infested Waters List through 2024 (85%; Table S1). We based our estimates of true invasion status on estimated rates of undocumented invasions in neighboring Wisconsin (Vander Zanden et al. 2024), where authors found the true invasion status was approximately four times higher than the documented status.

### Quantifying AIS Spread in Minnesota

#### Pattern analysis

From the Infested Waters List records, we calculated the numbers of new infestations per year per taxon through 2024. Then, we conducted trend analyses over the past four decades of these data (1985 to 2024) because most AIS were first confirmed in Minnesota in the late 1980s (Appendix S1: Table S1). First, we conducted breakpoint regression to test for significant changes in rates of new invasions over time using the *strucchange* R package (Zeileis et al. 2002). We selected the optimal number of breakpoints through comparison of model Bayesian Information Criteria. Second, to estimate continuous change in the rate of new infestations, we fit generalized additive models (GAMs) of the number of new infestations as a smoothed function of time with Normally distributed error using the R package *mgcv* (Wood 2015). We calculated model derivatives using the *gratia* package (Simpson and Singmann 2023). The parameter *k* (the upper limit on the number of basis functions used in the model, which determines “wiggliness”) was optimized using the function *gam.check*.

#### Basic Reproductive Number

Using an adaptation of the basic reproductive number (R_0_), we examined changes in AIS spread dynamics in Minnesota from 1984 to 2024. In epidemiological modeling, R_0_ is expressed as the product of the effective transmission rate and the duration of infectiousness. Conventionally, if R_0_ > 1, each infected individual, on average, produces more than one new infected individual, and the pathogen can invade the susceptible population and generate an epidemic (Anderson and May 1991). R_0_ = 1 creates conditions for endemicity, and an R_0_ < 1 leads to the infection fading out in the population. In the context of AIS, however, infested lakes often remain infested, so even relatively low invasion rates can yield an R_0_ value above 1. Because our focus was on evaluating the impact of prevention over time, versus estimating the average secondary infestations caused by a lake, we were able to overcome this limitation and extended this concept to estimate a time-varying reproductive number, R_t_, which allows us to assess changes in AIS spread among Minnesota’s lakes across successive time periods. We used the *EpiEstim* package in R (Cori et al. 2025) to estimate (R_t_). Because the timing of infestations between lakes varies, we used a parametric serial interval with a mean of 2 years and a standard deviation of 1 year, which represented an approximated time between reported infestations from a source lake to a recipient, reflecting the expectation that secondary spread become more likely to occur in years following establishment and population growth (Jones and Montz 2020). We estimated (R_t_) using a sliding estimation window of 7 years and reported a posterior means and the 95% credible intervals.

#### Simulation Modeling

The most compelling evidence to evaluate prevention effectiveness is counterfactual: how many otherwise-occurring invasions were prevented by the intervention? To answer this question, we used the model embedded in AIS Explorer (“Intervention Impact” application with code published alongside Angell et al. 2025). The network model simulates the movements of zebra mussels and starry stonewort between lakes using boater-movement and hydrological connections. Developed to aid local and statewide prevention planning, the model requires users to specify the effort and effectiveness of prevention efforts (i.e., boat inspections, decontamination, and education and outreach) at selected lakes (N. R. Angell et al. 2025). These manager-defined parameters then modulate the risk of future invasions spreading along edges of the network, but also provide the necessary dials to simulate AIS spread in Minnesota under different prevention effort scenarios.

To simulate what would have happened without as many statewide prevention efforts from 2016 to 2024, we used the Minnesota Infested Waters list to rewind the invasion status of zebra mussels and starry stonewort in Minnesota lakes in the model to 2015 levels: 173 zebra mussel infested lakes, and 1 starry stonewort infested lake. We then ran the model over 8 simulated years (corresponding to the time period 2016-2024), lightly tuning the underlying model parameters that determine invasive species spread likelihood until they produced the average rates of new infestations observed in that same time span (373 new zebra mussel and 29 new starry stonewort infestations) to serve as a baseline for comparison.

Then, to quantify intervention effectiveness, we obtained boat inspection and decontamination effort data from an ongoing project collating such data statewide (averaging 414,000 inspections annually across 691 Minnesota lakes from 2018-2023, conducted by over 60 local and tribal governments; Minnesota Department of Natural Resources 2025b). We combined these with conservative estimates for inspection and decontamination effectiveness: 70% from a recent study estimating these in Minnesota (N. Angell et al. 2024). Because we lacked data on education and outreach, we omitted such prevention in this analysis, which likely reduced our effectiveness estimates. We then functionally deducted these estimated interventions from the model to simulate what might have happened without them over 2016-2024 by increasing the per edge probability of infestation spread by a proportion between 0 and 1 corresponding to the intervention effort lost (for details see Angell et al. 2025). We conducted 250 simulations each with and without this prevention effort reduction to control for stochasticity and uncertainty in the model’s outputs of new zebra mussel and starry stonewort infestations.

## Results

### Is AIS prevention still warranted in Minnesota?

Using state invasion records and published estimates of lake true invasion rates and AIS suitability, we estimated Minnesota would currently be < 10% “saturated” with AIS (100% saturation is defined as all lakes being invaded with all species) in 11 of 15 scenarios and < 25% saturated in 14 of 15 scenarios (Figure 1). The only scenario suggesting > 25% saturation is one in which we know of only 25% of all current invasions but just 10% of lakes are invadable, a scenario past work suggests is unlikely (Muthukrishnan et al. 2018; Kanankege 2019).

**Figure 1.**
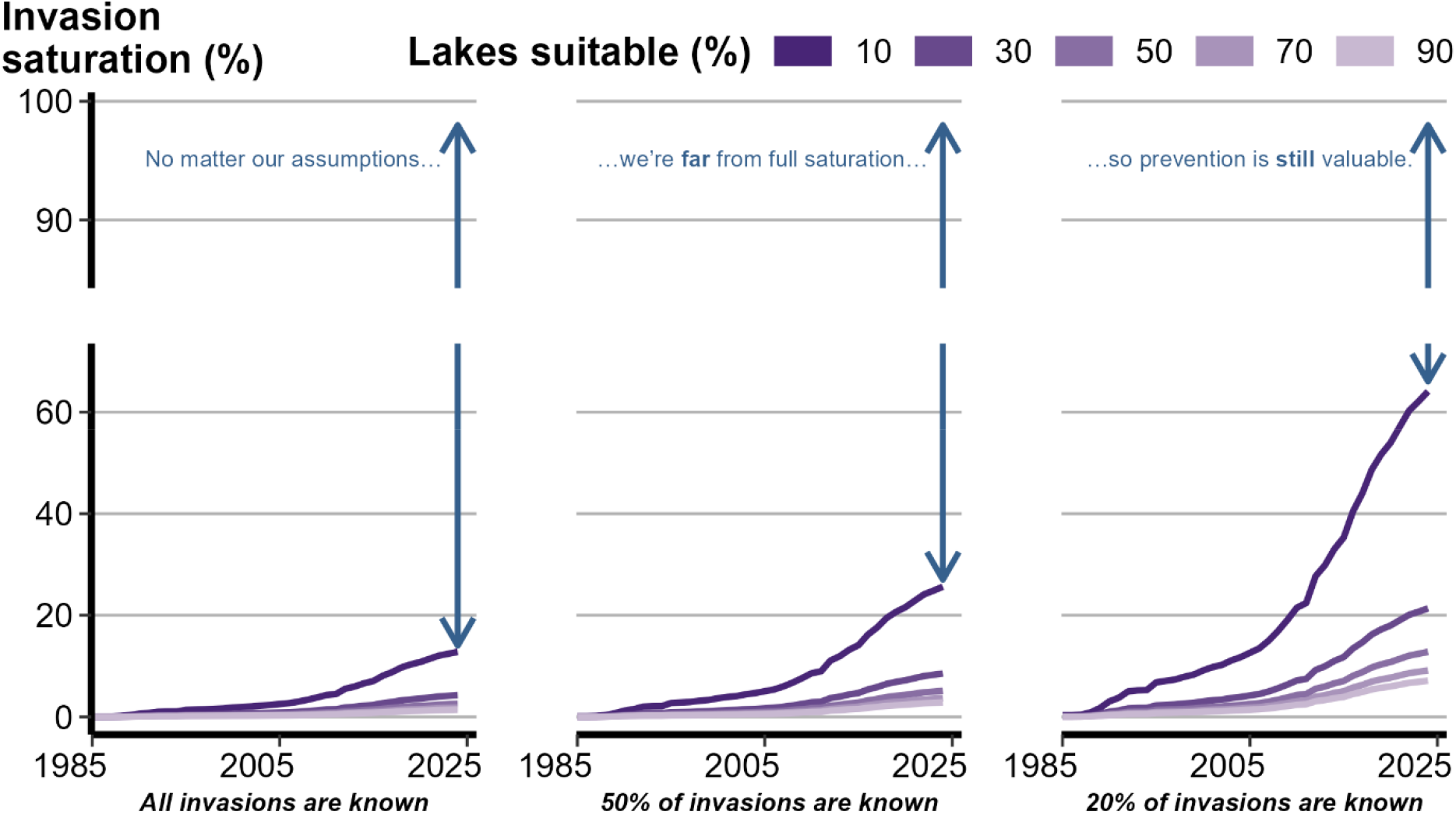
Estimates of invasion saturation in Minnesota lakes under different assumptions. 100% saturation was defined as all invadable lakes being invaded by all species. Panels are split by assumptions related to the % of true invasions known (i.e., documented in the MNDNR Infested Waters List). Assumptions related to the % of lakes suitable for invasion are represented by purple lines. Note the y-axis break between 60 and 90% saturation.

### Have rates of AIS spread in Minnesota changed?

While the cumulative number of AIS infestations has increased in Minnesota (Appendix S1: Figure S2), the rate of new infestations has been more dynamic. We detected a primary break in the rate of new infestations in 2015 (95% CI: 2011-2016) and a secondary break in 1992 (95% CI: 1992-1993; Figure 2a). The rate of new infestations accelerated from 1985-1992 (2.1 ± 0.62 SE infestations year^-2^) and accelerated at a slightly slower rate from 1992-2014 (1.6 ± 0.18 SE infestations year^-2^) before decelerating from 2016-2024 (−4.0 ± 1.2 SE infestations year^-2^). The rate of new infestations estimated with the fitted GAM was relatively stable through 2001, accelerated from 2002 to 2014 (reaching a peak rate of 3.4 ± 0.52 SE infestations year^-2^ in 2009), crossed 0 infestations year^-2^ between 2017 and 2018, and significantly decelerated from 2022 to 2024 (−2.0 ± 0.95 SE infestations year^-2^ in 2024; Figure 2c).

**Figure 2.**
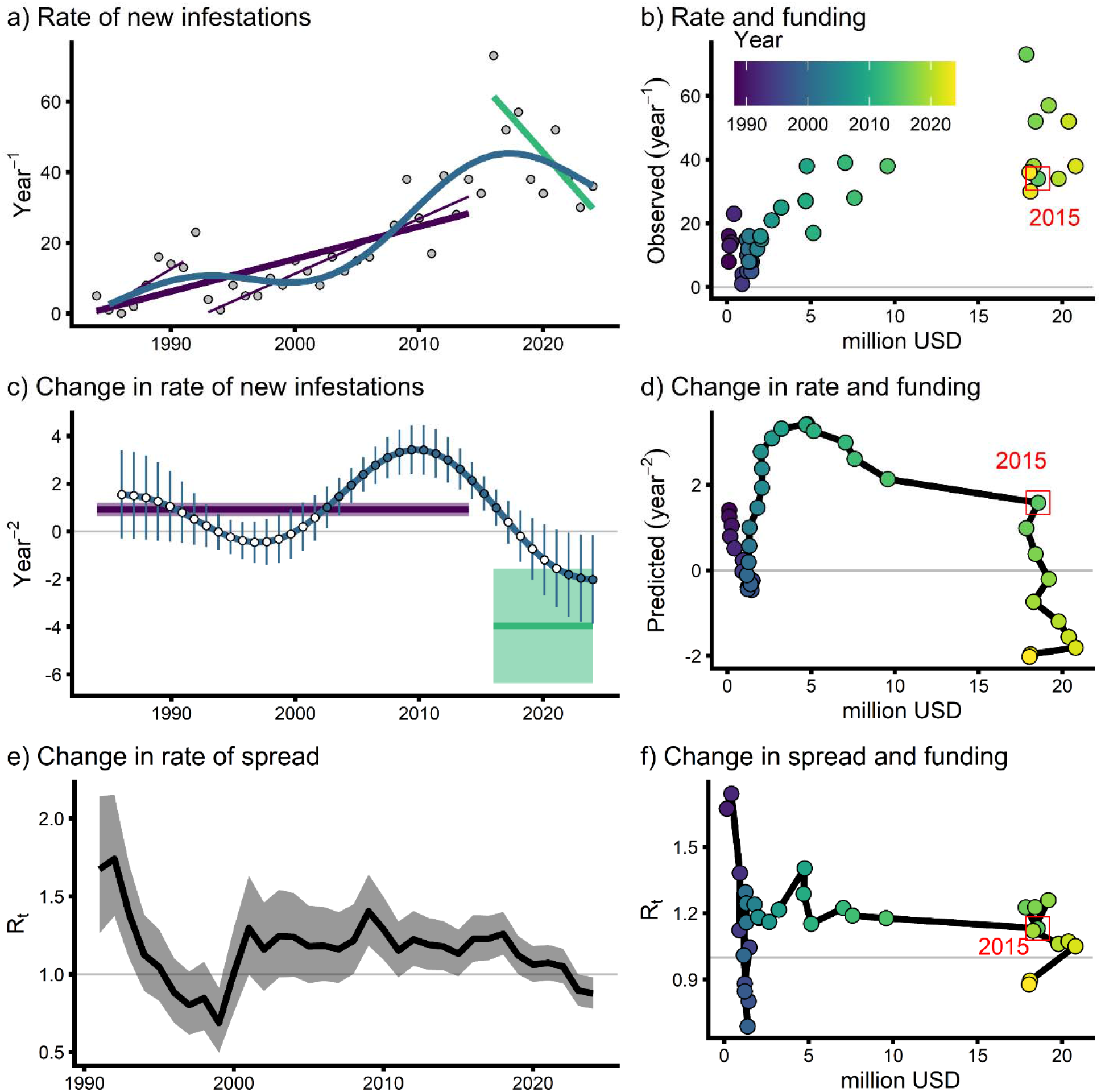
Tracking AIS infestation status (a, c, e) and funding (b, d, f) in Minnesota over time. a) The observed rate of new infestations (infestations year^-1^) is plotted in grey circles with fitted trend lines for the breakpoint analysis (1985-1992 and 1992-2014 in thin purple lines, 1985-2014 in a thicker purple line, 2016-2024 in a thicker green line) and shows accelerating infestations from 1985 to 2014 and decelerating infestations from 2016-2024. The trend fitted GAM model is plotted with a blue line and tracks with the breakpoint trend lines. b) Observed rate of new infestations is plotted with invasive species prevention and management funding to the DNR and Counties in Minnesota (Appendix S1: Figure S1). Brighter green and yellow indicates more recent years with 2015 highlighted with a red box and labeled with red text. c) Change in the rate of infestations (infestations year^-2^) is represented by the first order derivative from the fitted GAM model in panel “a” (blue line and circles with point-level 95% Confidence Intervals and significant periods of change plotted with filled circles) and slopes from the linear trends in panel “a” (purple and green line with 2 SE around the slope). Both pattern analysis approaches show decelerating infestation rates in the past decade. d) GAM-predicted change in the rate of infestations is plotted against funding as in panel “b”. e) Estimated reproductive number (R_t_) is plotted in a black line with 95% Confidence Interval in shaded grey. f) Reproductive number is plotted against funding as in panel “b”.

Decelerating invasion rates occurred alongside increasing AIS management and prevention funding (Figure 2b, d). Total State funding to the DNR and to Counties for invasive species management and prevention (Appendix S1: Figure S1) doubled in 2015, which corresponds to the first break in the breakpoint analysis and preceded when the derivative of the fitted GAM crossed 0 infestations year^-2^.

We found that the time-varying reproductive number, R_t_, declined over the study period, suggesting a long-term reduction in the relative spread intensity across lakes in Minnesota (Figure 2e). In the early 1990s, R_t_ values were high (∼1.5-2.0). R_t_ dropped sharply through the late 1990s, reaching values below 1, suggesting a shift towards lower spread. Between approximately 2000 and 2015, R_t_ stabilized at 1.0-1.3, with moderate fluctuations, indicating periods of sustained spread interspersed with relative declines. After 2015, R_t_ values exhibited a gradual downward trend, reaching values below 1.0 in the early 2020s. This suggests a reduction in the number of new infestations relative to the number of previously infested lakes, consistent with lower relative spread and effective prevention. Overall, the trend indicates a transition from early rapid expansion toward lower relative rates of spread, with recent R_t_ values suggesting that secondary infestations are occurring at a lower rate.

### How many invasions have prevention efforts prevented?

Using model simulations grounded in statewide real-world boater movement data and management effectiveness estimates, we estimated that prevention efforts from 2016-2024 may have prevented an average of 71 new zebra mussel invasions (354 mean infestations with prevention compared to 425 without, a 20% difference) and 16 new starry stonewort invasions (30 mean infestations with prevention compared to 46 without, a 53% difference) (Figure 3).

**Figure 3.**
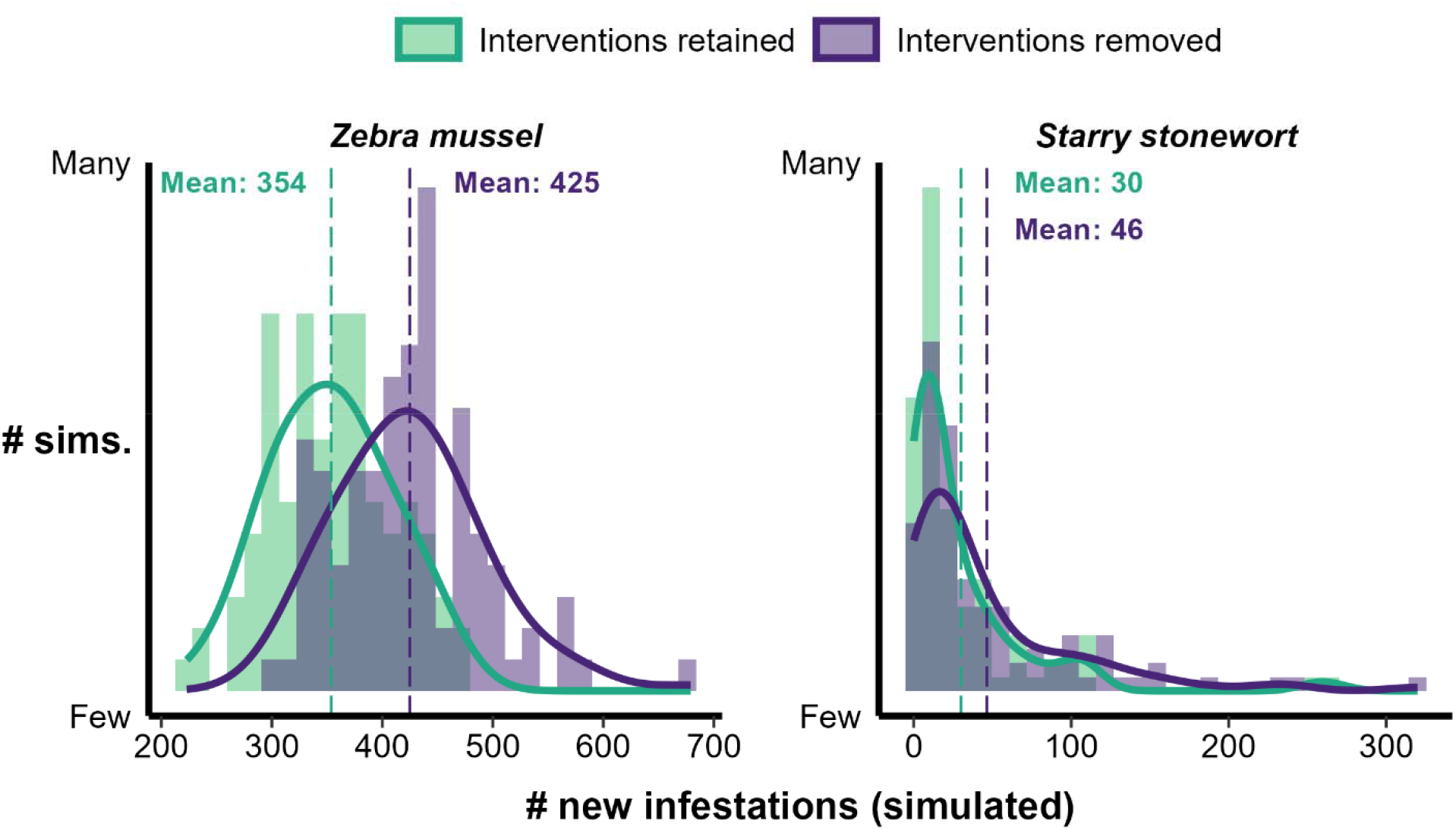
Simulation-model-derived estimates of numbers of new zebra mussel and starry stonewort infestations with all prevention interventions retained (green) versus with many known interventions removed (dark purple), over 250 simulations. Mean “interventions retained” values align with observed rates of new documented infestations during the time period of 2016-2024.

## Discussion

Our analyses provide concurrent evidence that AIS invasion rates have slowed since 2015 in Minnesota, coinciding with substantial increases in statewide investment in AIS management. This shift is not explained by saturation of suitable habitat, as Minnesota’s lakes likely remain far from fully invaded based on known and literature-based estimates of true infestation status and lake suitability. Our pattern analyses, saturation results, R_t_ estimates, and simulation modeling outputs suggest that prevention efforts have helped shift invasion dynamics from an early, expansion phase marked by unregulated exponential growth towards a slower, saturation-like phase (Arim et al. 2006), reinforcing the importance of sustained prevention effort to protect uninvaded lakes.

Evaluating the effectiveness of prevention requires multiple analyses because no one method fully captures the complexity of invasion processes (Epanchin-Niell 2017). By integrating several approaches, we arrived at a consistent finding: AIS spread has slowed during a period of significant prevention investment. It is essential to note that the time-varying reproductive number (R_t_) and the simulation model are grounded in shared concepts of spread dynamics, rooted in epidemic theory and centered around the idea that each infested lake can generate new infestations within the system. In this study, R_t_ represents an empirical measure of system-wide spread intensity. In contrast, the simulation model provides a mechanistic representation of that process, in which the invasion intensity emerges from lake invasion probabilities informed by historical infestations, boater movements, and hydrologic connections (Kinsley et al. 2024; N. R. Angell et al. 2025). Therefore, their agreement is validating, as they capture complementary processes underlying the same spread dynamics.

Investigating the effectiveness of invasive species prevention has been a longstanding challenge in invasion science. While our results provide evidence in support of prevention, some challenges and uncertainties remain. First, the MNDNR Infested Waters List only tracks a subset of AIS in Minnesota. Important but unlisted AIS include two widespread, long-established species: curly-leaf pondweed (*Potamogeton crispus*) and common carp (*Cyprinus carpio*). However, management of these species often focuses on control rather than preventing/slowing spread (Sharov and Liebhold 1998). Second, in our simulation model, we could not subtract 1) interventions not documented to the MNDNR via voluntary inspection reporting, nor 2) interventions related to education and outreach efforts. These are sizable omissions. Coupled with our choice of average effectiveness values from Angell et al. (2024), we suspect we have underestimated the number of new invasions prevented over the past decade. Granted, we cannot know if all our model’s assumptions are valid, but we were encouraged that, with modest tuning, the model produced results centered on the actual new infestation rates documented during the same time frame.

Our results provide multi-faceted support for invasive species prevention and were made possible by a long-term, statewide commitment to AIS management, documentation, monitoring, and research. Most notably, the clear inflection points in the spread rate of AIS in Minnesota, concomitant with a large increase in AIS management support, suggests these investments have been worthwhile. While future funding and the status of AIS in Minnesota remain unpredictable, especially with an expected reduction of $5 million in State funding to counties, we provided evidence that supports prevention as a tool to protect ecosystems and economies from the harmful effects of invasion.

## Acknowledgements

This work was funded by the Minnesota Environment and Natural Resources Trust Fund and the Minnesota Agricultural Experiment Station. We thank Molly Tilsen for her significant contributions to data curation, specifically her extensive work in compiling and cleaning the unpublished MNDNR watercraft inspection records used in this study. We would like to thank the countless State, local, and tribal natural resource managers and the public engaged in AIS prevention in Minnesota. These analyses, and the outcomes observed, would not have been possible without their dedication to AIS management, monitoring, and research.

## Author Contributions

JW led the manuscript drafting process. All authors contributed to the study conception and design, data analysis, and manuscript drafting and revisions. All authors read and approved the final manuscript.

