## Supplemental Information for "Slowed aquatic invasion coincides with intensified research and prevention in Minnesota, USA"

*bioRxiv – Supplementary Information*

### Appendix S1

“Slowed aquatic invasion coincides with intensified research and prevention in Minnesota, USA”

Jake R. Walsh, Alex W. Bajcz, Amy C. Kinsley, Nicholas B. D. Phelps

#### Contents:

Invasive Species List in the MNDNR Infested Waters List (Table S1)

Minnesota AIS Prevention Funding (Figure S1)

Cumulative Infestation Trends Over Time (Figure S2)

Appendix S1 References

### Invasive Species Listed in the MNDNR Infested Waters List

Table S1. Invasive species listed in the MNDNR Infested Waters List (N = number of lakes infested with a species through 2024, Confirmed = first year confirmed in a Minnesota water body).

| Scientific Name | Species Name | N | Confirmed |
| --- | --- | --- | --- |
| Eurasian Watermilfoil | *Myriophyllum spicatum* | 389 | 1987 |
| Zebra Mussel | *Dreissena polymorpha* | 340 | 1989 |
| Flowering Rush | *Butomus umbellatus* | 45 | ≤1984 |
| Spiny Water Flea | *Bythotrephes longimanus* | 30 | 1987 |
| Starry Stonewort | *Nitellopsis obtusa* | 30 | 2015 |
| Faucet Snail | *Bithynia tentaculata* | 29 | 2008 |
| Brittle Naiad | *Najas minor* | 10 | 2003 |
| Bighead Carp | *Hypophthalmichthys nobilis* | 7 | 2003 |
| Grass Carp | *Ctenopharyngodon idella* | 6 | 2008 |
| Silver Carp | *Hypophthalmichthys molitrix* | 6 | 2008 |
| Red Swamp Crayfish | *Procambarus clarkii* | 1 | 2016 |
| Viral Hemorrhagic Septicemia Virus |  | 1 | 2010 |
|  | Total | 894 |  |

#### Minnesota AIS Prevention Funding

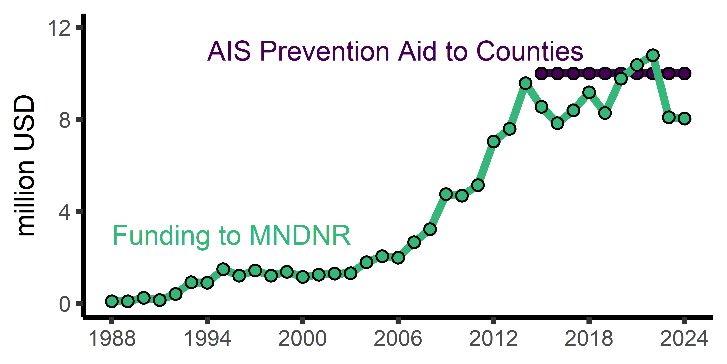

Figure S1. Two examples of Statewide funding for invasive species management from 1988-2024 (funding in millions of US dollars to Minnesota counties in dark purple and the MN Department of Natural Resources in green). Funding to the MNDNR includes ~$250,000 per year for terrestrial invasive species management. MNDNR funding in 2016 is interpolated (empty circle).

The MNDNR Invasive Species Program is funded primarily through annual State general-fund appropriations, a boater registration surcharge, a non-resident fishing license fee, and federal grants. In addition to occasional one-time appropriations, increases to funding since 1988 have been due to an increase in the boater registration surcharge (1.00 USD to 5.00 USD from 1990-1994 and to 10.60 USD in 2020 for three-year registrations), an increase in the non-resident fishing license fee (established at 2.00 USD in 2008 and increased to 5.00 USD in 2013), and the addition of federal grants to the budget (including ~1 million USD each year since 2010 from the US Fish and Wildlife Service to implement the Minnesota Management Plan for Invasive Species).

### Cumulative Infestation Trends Over Time

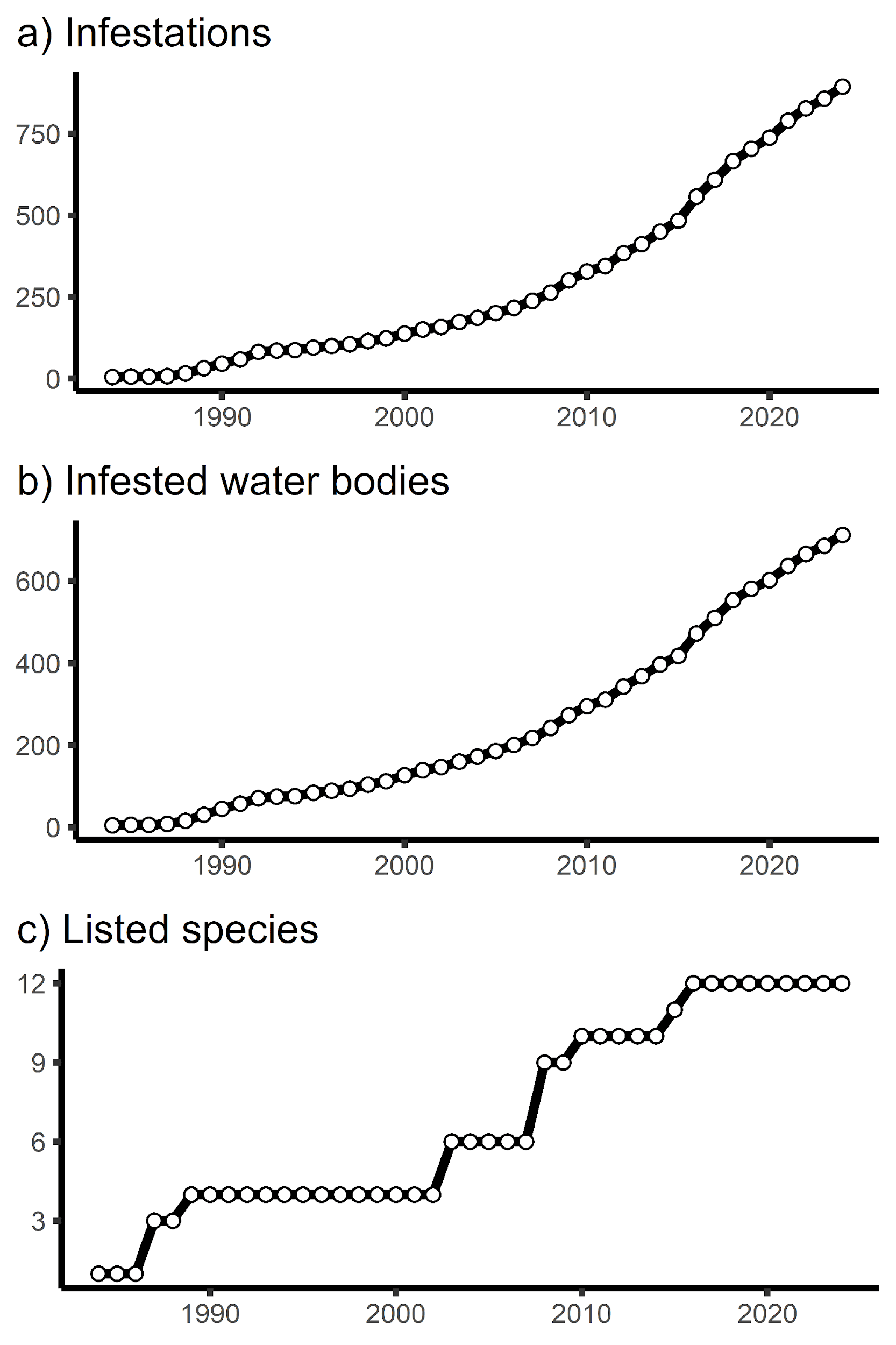

Figure S2. Cumulative aquatic invasion records for a) infestations, b) infested water bodies, and c) listed AIS.
